# Towards a systematic characterization of protein complex function: a natural language processing and machine-learning framework

**DOI:** 10.1101/2021.02.24.432789

**Authors:** Varun S. Sharma, Andrea Fossati, Rodolfo Ciuffa, Marija Buljan, Evan G. Williams, Zhen Chen, Wenguang Shao, Patrick G.A. Pedrioli, Anthony W. Purcell, María Rodríguez Martínez, Jiangning Song, Matteo Manica, Ruedi Aebersold, Chen Li

## Abstract

It is a general assumption of molecular biology that the ensemble of expressed molecules, their activities and interactions determine biological processes, cellular states and phenotypes. Quantitative abundance of transcripts, proteins and metabolites are now routinely measured with considerable depth via an array of “OMICS” technologies, and recently a number of methods have also been introduced for the parallel analysis of the abundance, subunit composition and cell state specific changes of protein complexes. In comparison to the measurement of the molecular entities in a cell, the determination of their function remains experimentally challenging and labor-intensive. This holds particularly true for determining the function of protein complexes, which constitute the core functional assemblies of the cell. Therefore, the tremendous progress in multi-layer molecular profiling has been slow to translate into increased functional understanding of biological processes, cellular states and phenotypes. In this study we describe PCfun, a computational framework for the systematic annotation of protein complex function using Gene Ontology (GO) terms. This work is built upon the use of word embedding— natural language text embedded into continuous vector space that preserves semantic relationships— generated from the machine reading of 1 million open access PubMed Central articles. PCfun leverages the embedding for rapid annotation of protein complex function by integrating two approaches: (1) an unsupervised approach that obtains the nearest neighbor (NN) GO term word vectors for a protein complex query vector, and (2) a supervised approach using Random Forest (RF) models trained specifically for recovering the GO terms of protein complex queries described in the CORUM protein complex database. PCfun consolidates both approaches by performing the statistical test for the enrichment of the top NN GO terms within the child terms of the predicted GO terms by RF models. Thus, PCfun amalgamates information learned from the gold-standard protein-complex database, CORUM, with the unbiased predictions obtained directly from the word embedding, thereby enabling PCfun to identify the potential functions of putative protein complexes. The documentation and examples of the PCfun package are available at https://github.com/sharmavaruns/PCfun. We anticipate that PCfun will serve as a useful tool and novel paradigm for the large-scale characterization of protein complex function.

## INTRODUCTION

Proteins are known to catalyze and control the vast majority of the reactions of cellular biochemistry (Aebersold and Mann, 2016). Frequently they exert their function only if they stably interact in precise stoichiometric ratios with other proteins in the form of complex macromolecular structures, a notion that has been encapsulated in the term “modular cell biology” (Hartwell et al., 1999). With the advent of high throughput ‘OMICS’ technologies for the study of complex biological systems, it is now possible to accurately quantify and identify different types of biologically relevant molecules across various conditions. However, determining the biological functions and phenotypes of these assemblies has remained challenging and requires a functional understanding of the molecular functions of its components and associations. Detailed biochemical and cell biological studies have identified the composition and even the atomic structures of numerous protein complexes with well-defined roles in a variety of fundamental biological processes (Hewick et al., 2003), such as in their participation in transcriptional regulation (Aranda et al., 2015; Simonis et al., 2004; Tan et al., 2007; Webb and Westhead, 2009), cell cycle control (Becher et al., 2018; Chen et al., 2019; D’Avino et al., 2009) and signal transduction (Pawson and Nash, 2000; Rebois and Hebert, 2003). Protein complexes can, therefore, be considered essential agents and indicators of cellular functionality.

Recent technical advances, particularly in mass spectrometry (MS) - based proteomics have greatly enhanced our capacity to determine the composition, stoichiometry and abundance of known protein complexes and to identify new entities. These methodologies also support the systematic identification of compositional or quantitative changes in complexes as a function of cellular state. These include methods such as BF-MS (Biochemical Fractionation Mass Spectrometry) (Carlson et al., 2019; Heusel et al., 2019; Heusel et al., 2020; Rosenberger et al., 2020; Stacey et al., 2017; Szklarczyk et al., 2019), Affinity Purification MS (AP-MS), Cross-Linking MS (XL-MS) (Leitner et al., 2016; Leitner et al., 2012; Liu et al., 2015) and limited proteolysis (LiP) (Schopper et al., 2017) and thermal proteome profiling (TTP) (Mateus et al., 2020). Compared to the experimental detection of new protein complexes, the determination of their biochemical or cellular function has significantly lagged behind because experimentation with specific complexes is highly challenging. Given the challenge of characterizing the function of protein complexes, hypotheses regarding the functional roles in which a newly discovered protein complex participates are typically generated by careful manual review of prior literature.

The standard approach to manual literature review for identifying the putative function of a protein complex consists of the search for publications and database entries about the individual protein subunits and followed by the consolidation of the retrieved information. However, this manual curation presents several limitations that cam make it highly inefficient and highly biased. First, exhaustive literature curation for all proteins belonging to even a single complex can easily become prohibitively time-consuming due to the sheer volume of publications required to parse through. Given that the manual curation for retrieving high-confidence functional annotations of a single protein complex can be extremely laborious, performing such annotation on dozens or hundreds of novel entities discovered in large scale complex centric proteomic fractionation experiments quickly becomes prohibitive. Second, although protein complex databases, such as CORUM (Giurgiu et al., 2019) and Complex Portal (Meldal et al., 2019) offer experimentally and manually validated functional annotations for better-studied protein complexes (i.e. the ground truth), the literature-based functional annotations for the same protein complexes can be highly dissimilar across different databases. Third, some proteins are multifunctional and may have unique roles in different protein complexes, thereby highlighting that the function of protein complexes is not simply the aggregate of their subunits’ functions (Jeffery, 2015; Matalon et al., 2014; Nakabayashi et al., 2014). As a case in point, we conducted a preliminary examination of the Gene Ontology (GO) terms annotated for whole protein complexes in the CORUM database compared to each individual subunit’s GO term annotations in the QuickGO database (Binns et al., 2009). The results showed that 2155 (61.4%), 319 (9.1%), and 169 (4.8%) heteromeric protein complexes in CORUM contained at least one novel biological process, molecular function, or cellular component term that was not annotated for any individual subunit’s QuickGO entry, respectively. In other words, certain proteins may participate in emergent functionality when assembled in a macromolecular complex that would be non-obvious based on the known functions of the individual protein complex’s subunits. We therefore argue that it is of great importance to employ computational techniques that can assist large-scale predictions of protein complex functions and provide useful insights and guidance for the follow-up functional characterization experiments.

Given that the nature of information encoded in natural language-based functional descriptions of protein complexes is fundamentally unstructured, tit is challenging for traditional bioinformatic and data mining approaches to meaningfully distill the information from specific publications. However, computational methods from text-mining — the field concerned with computationally extracting information from unstructured natural language text— have been also successfully applied to a variety of biomedical problems and provide a promising avenue to address our task. These include extraction of protein-protein relations and functions (Islamaj Dogan et al., 2019; Li et al., 2019b; Manica et al., 2019; Subramani et al., 2015; Yu et al., 2018), determination of protein structure (Gaizauskas et al., 2003), protein localization (Cejuela et al., 2018), and gene-disease relationships (Pletscher-Frankild et al., 2015). However, to the best of our knowledge, no computational tool designed specifically for annotating the functions of protein complexes has been described to date.

To address this dearth in direct functional annotation methods for protein complexes, in this work we integrate text-mining and machine-learning techniques into a hybrid computational framework, termed PCfun, that can be applied to large scale complex-centric proteome experiments for predicting the function of protein complexes. At a high level, PCfun is developed based upon word embedding generated from the machine reading of >1 million open access PubMed Central (PMC) articles, whereby both unsupervised and supervised machine learning algorithms were used to generate two separate lists of predicted functional Gene Ontology (GO; biological process, molecular function and cellular component) terms for a queried protein complex. Following, a supervised machine-learning model trained on the associations between protein complexes and their GO terms documented in the CORUM database was used to predict a second list of candidate functional terms. Hence, the unsupervised candidate list provides functional predictions solely based upon the word vector relationships observed within the embedding that are unbiased to protein complex-function associations while the supervised candidate list tailors the annotations to relationships similar to the CORUM database. In order to leverage the insights provided by both approaches we attempted to consolidate the two lists by leveraging the hierarchical structure of the Gene Ontology by testing for enrichment of certain supervised terms within the unsupervised list. An adapted leave-one-out cross-validation scheme was used to test the system’s performance and suggested that PCfun achieved outstanding prediction performance with AUC values of 0.895, 0.927, and 0.957 for biological process, molecular function, and cellular component terms, respectively. In addition, we compared the prediction outcomes by PCfun and the GO annotations from the Complex Portal database (Meldal et al., 2019) using protein complexes not documented in the CORUM database. For the biological function and cellular component categories, PCfun predicted similar (i.e. semantic similarity >= 0.5) GO terms that covered more than half of the Complex Portal’s ground-truth annotations for 52.8% and 69.7% of the protein complexes in the biological process and cellular component categories respectively. In contrast, the molecular function category only achieved 12.9% coverage, which might be explained by the lowest similarity between the CORUM and Complex Portal annotated GO terms for this category. Taken together, we anticipate that PCfun will serve as an accurate annotation tool for protein complex function and increase our better understanding of the functional roles of protein complexes in biological systems.

## RESULTS

### The Architecture of PCfun for Predicting the Function of Protein Complexes

PCfun development consisted of two main steps. The first step, as shown in **Figure 1A**, was based on the building of the word embedding itself. Approximately 1 million open access articles were downloaded from the PubMed Central Repository and their full texts were processed (STAR Methods) as described in Manica *et al.* (Manica et al., 2019) to populate a text corpus. After consolidation of the text corpus, the fastText implementation of the skip-gram context prediction embedding algorithm was employed onto the text corpus (STAR Methods) to construct a word-embedding representation of the text: 500-dimensional, continuous real-valued vector representations based on the subwords extracted from the corpus. These word vectors were constructed for character *n*-grams allowing for the creation of a word vector for any natural language query (see **Figure 2A** for a graphic illustration of the word vectors). Using this property of character *n*-gram embeddings, we next extracted sub-embeddings consisting of all protein complex and GO term (split into biological process, molecular function, and cellular component classes) queries. As a result of this first step (**Figure 1A**) we obtained five-word vector sub-embeddings corresponding to protein complex queries (sub-embedding for each naming scheme used for a protein complex name: canonical *vs.* subunit name) and GO term queries (three sub-embeddings corresponding to biological process, molecular function, and cellular component terms).

**Figure 1.**
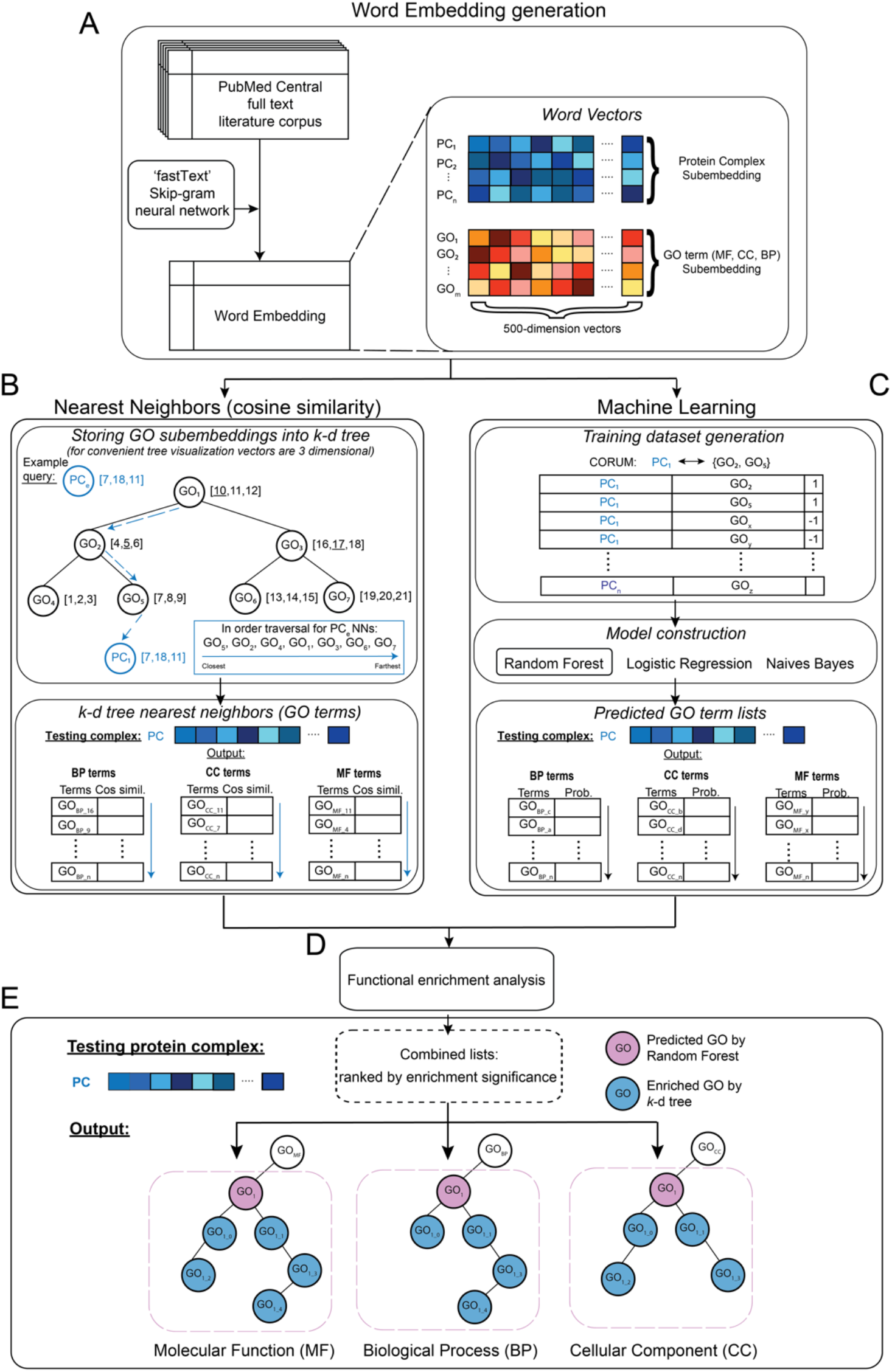
The overall framework of PCfun. (A) A word-embedding containing a 500-dimentional vector for each word was first generated based on the open-access full-text articles and their abstracts using the ‘fastText’ with a skip-gram model. Based on the word-embedding, two machine-learning algorithms were used, including (B) a *k*-d tree for nearest-neighbor search, and (C) a supervised RF model for PC association with MF, BP and CC, respectively. A simplified *k*-d tree example is shown in the top panel of (B). To combine the outputs of the two models, GO terms enrichment analysis (D) was performed. PCfun utilizes the enrichment analysis and GO DAG structure (E) to represent and visualize the predicted GO terms for a given protein complex. The testing protein complex (i.e. ‘PC’) is given to illustrate the usage of PCfun.

**Figure 2.**
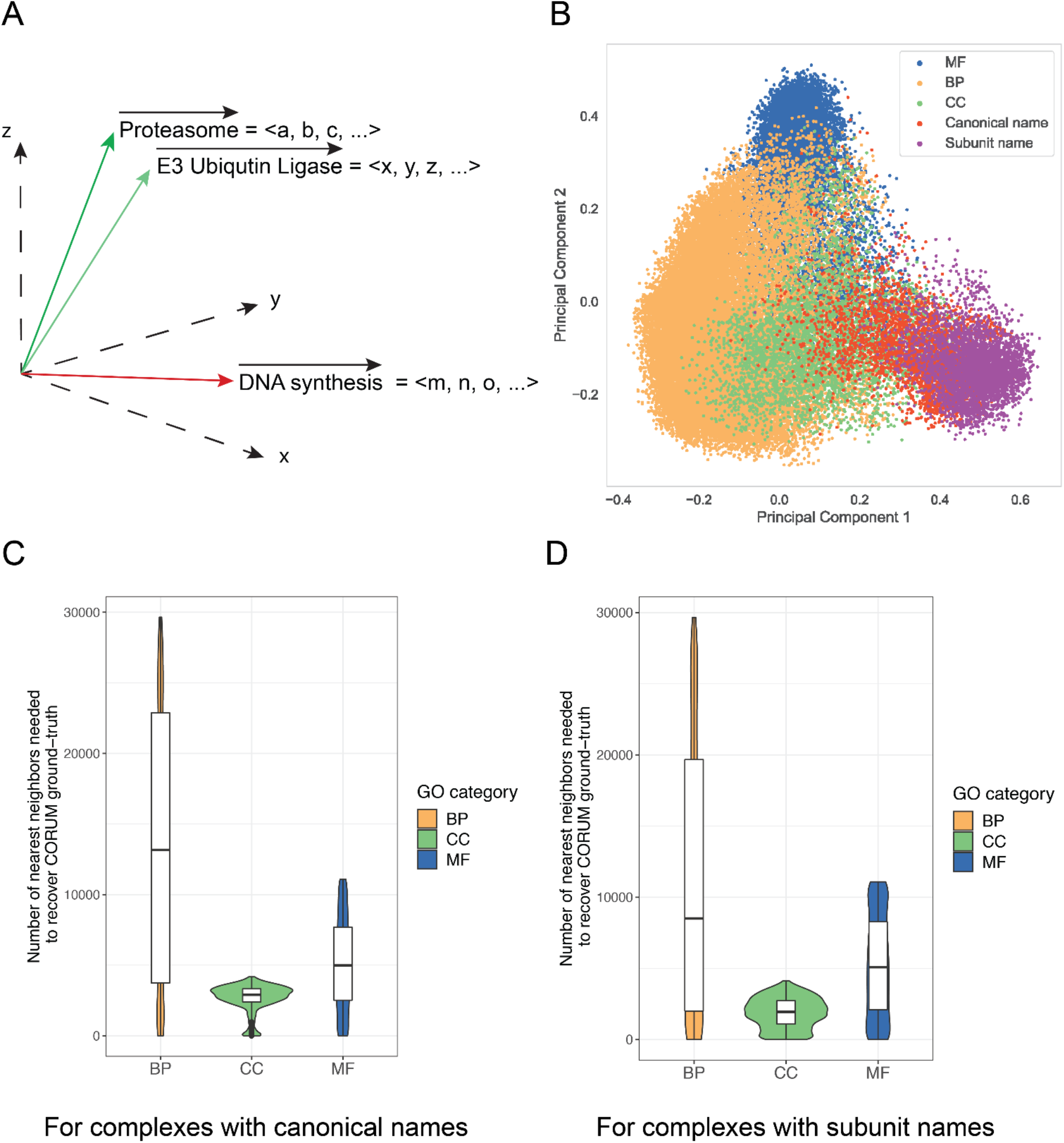
Using the word-embedding and *k*-d tree to shortlist GO terms for protein complexes. (A) A graphical illustration of the vectors of phrases ‘Proteasome’, ‘E3 Ubiquitin Ligase’ and ‘DNA synthesis’ in a 3D space using simplified vector representation, for example, < *a, b, c,* … > denotes the numerical word vector for the natural language query ‘Proteasome’. (B) The principal component analysis (PCA) results of different types of sub-embeddings, including molecular function, biological process, cellular component, protein complexes with canonical names, and subunit names. (C) The numbers of nearest neighbors required from the *k*-d tree search outputs for protein complexes using canonical names to cover the CORUM ground-truth. (D) The numbers of nearest neighbors required from the *k*-d tree search outputs for protein complexes using subunit names to cover the CORUM ground-truth.

The next step was then to construct the models for functional annotation of protein complex queries. As depicted in **Figures 1B** and **C**, we employed two strategies capable of returning ranked protein complex – GO associations: (i) an unsupervised nearest-neighbor approach illustrated in **Figure 1B**, and (ii) a supervised machine learning approach displayed in **Figure 1C**. The first algorithm is agnostic to the question of functional annotation of protein complexes and was based solely upon contextual relationships. In contrast, the second algorithm is a tailored approach trained specifically to recover functional terms for a protein complex query. The rationale for using two distinct approaches is that they are likely to produce complementary, and potentially, if combined, more informative outputs.

As seen in **Figure 1B**, we built a *k*-d tree (*k*-dimensional tree) (Freidman et al., 1977), a space-partitioning structure for storing the sub-embeddings’ vectors of GO terms to enable rapid application of a nearest neighbor algorithm that was able to shortlist GO terms ranked by cosine similarity between the queried protein complex vector and each word vector for a GO term, for recovering CORUM’s ground-truth annotations of each protein complex. The supervised machine-learning models (**Figure 1C**), on the other hand, learned from the experimentally verified protein complex – GO term associations and were therefore able to accurately cover these ground-truth in the CORUM database. We constructed and evaluated four widely applied machine-learning algorithms: RF (Breiman, 2001), Logistic Regression (LR) (Lecessie and Vanhouwelingen, 1992; Yu et al., 2011), and Naïve Bayes (NB) classifier a with Gaussian distribution and a Bernoulli distribution (Zhang, 2004) classifiers. A ranked list of GO term annotations was generated by both the unsupervised *k*-d tree algorithm (**Figure 1B**) and the supervised machine learning models (**Figure 1C**).

Finally, to combine the prediction outcomes from the RF and the *k*-d tree, a hypergeometric test was conducted to test for the functional enrichment of *k*-d tree terms within the child nodes of each RF predicted term (**Figure 1D**). A visualization of a GO direct acyclic graph (DAG) structure for functionally enriched predicted GO terms was performed to represent the contextual information of predicted GO terms of biological process, molecular function and cellular component, respectively (**Figure 1E**). Given a protein complex of interest, PCfun first applies the two models to generate two prediction lists using *k*-d tree and RF, and then visualizes the prediction outcome via the functional enrichment analysis and the GO DAG structure (STAR Methods).

### Benchmarking the prediction performance of PCfun

We systematically evaluated the prediction performance of PCfun. We first separately assessed the predictive ability of the unsupervised *k*-d tree and the RF model, for annotating protein complex functions. Further, we independently compared the prediction outputs from the enrichment analysis of PCfun with the functional annotations documented in the Complex Portal database (Meldal et al., 2019).

#### The word-embedding and k-d tree facilitated the ranking of potential GO terms for protein complexes

A useful property of word-embeddings is that words with related semantic meanings have corresponding word vectors that exist closer to each other in the word vector space - as measured by the cosine similarity (i.e. same orientation) – compared to words that have very different meanings. Therefore, one can find words with a similar meaning to an input query word by simply finding the nearest neighbors of the input query word vector. To aid in rapid nearest-neighbor calculations for these large sub-embeddings, we stored each sub-embedding into a *k*-d tree, allowing us to efficiently retrieve word vectors that were similar to the input query vector. To evaluate the quality of the word-embeddings, we performed principal component analysis (PCA) of the word vectors for each extracted sub-embedding of different types, including biological process vectors, molecular function vectors, cellular component vectors, and the protein complex vectors with the two naming schemes (STAR Methods). **Figure 2B** demonstrates that the sub-embeddings’ word vectors of each type are well clustered, indicating the reliable quality of the word-embedding.

We measured the ability of each GO term class sub-embedding to recover the ground-truth functional annotations for a protein complex from CORUM by recording the number of nearest neighbors (ranked by their cosine similarity) required to recover 100% of the ground-truth functional annotations for an input protein complex query. We hypothesized that the results might change depending on the name used to represent a protein complex. Additionally, considering that de novo detected protein complexes will not be characterized with an accepted name, we proposed the subunit naming scheme for a protein complex that would still allow for the functional annotation of even newly identified protein complexes by PCfun. Therefore, we tested the two protein complex naming schemes’ sub-embeddings (STAR Methods). **Figures 2C, D** indicate that the sub-embeddings required on average 13487, 5119, 2692, and 11044, 5214, 1894 nearest neighbors to recover the ground truth for biological process, molecular function, and cellular component categories using the canonical names and subunits names, respectively. It is evident that in order to recover CORUM’s ground-truth annotations, a large number of nearest neighbors are required. We therefore subsequently performed manual literature search based on the top nearest neighbor GO terms for certain protein complexes and observed that the predicted GO terms were actually still quite informative and were recovering known biological knowledge.

We chose a representative example protein complex, namely “SMAD2-SMAD4-FAST1-TGIF-HDAC1 complex, TGF (beta) induced”, for a manual literature review comparison to the *k*-d tree nearest-neighbor results. We used the subunit naming scheme (“smad4 tgif1 smad2 hdac1 foxh1”) for generation of its corresponding word vector and then queried the vector into the biological function, molecular function and cellular component sub-embedding *k*-d trees. While the *k*-d trees required 27400, 2182 and 990 nearest neighbor terms in the biological function, molecular function and cellular component trees, respectively, to recover the six CORUM annotated GO terms (DNA topological change; negative regulation of transcription, DNA-templated; DNA binding; transforming growth factor beta receptor signaling pathway; chromosome organization; nucleus) for this protein complex, the top returned *k*-d tree nearest neighbors (**Supplemental Table S1**) still provided relevant GO terms that had related biological meanings. For example, the top 10 nearest neighbors for the biological function category are terms all related to the TGFβ or bone morphogenic protein response. According to Massague *et al* (Massague et al., 2005), the SMAD proteins accumulate in the nucleus to execute transcriptional control in response to TGFβ signal transduction and may be co-activated or co-repressed by various DNA-binding co-factors. We observed that “negative regulation of Smad protein signal transduction” was the 8th nearest neighbor term for the queried protein complex vector, which recovers the role of the co-repressor activity of HDAC1 and TGIF that act to repress the transcriptional control of the activated and nuclear localized SMAD2:SMAD4 subcomplex (Liberati et al., 1999; Wicks et al., 2000). The ranking of GO terms by the *k*-d tree for this protein complex is listed in **Supplementary Files 1-3**. In summary, the top-nearest neighbor results of the *k*-d tree do provide insights into the relevant biology, but demonstrate poor ability to recover CORUM’s ground-truth annotations. This suggests the necessity of building supervised-learning models to systematically and statistically improve the predictive outcomes.

#### Supervised machine-learning models greatly improved the performance of ground-truth recovery of GO terms in CORUM

In order to improve the performance of ground-truth recovery of CORUM, we implemented supervised machine-learning classifiers based on the word vectors for a ‘protein complex-GO association’ pair (termed ‘PC-GO’). In our study, the annotated association of a PC-GO term was regarded as a positive sample, whereas randomly sampled synthetic pairs of a protein complex and other GO terms that were not associated in CORUM were regarded as negative ones. As the negative samples significantly outnumbered the positive samples in the resulting datasets, we generated five different training datasets with randomly selected negative samples and all positives for each protein complex to ensure an equal distribution of positive and negative samples for training the classifier (STAR Methods). This process was conducted for both naming schemes. With the training datasets, we assessed the performance of three machine-learning classification algorithms, including RF, LR, and NB with a Gaussian or Bernoulli prior (NB_Gauss or NB_Bernoulli, respectively), through the adapted ‘protein complex’-leave-one-out cross-validation strategy using standard performance measures (STAR Methods).

Across these classifiers, RF consistently performed the best as measured by all performance metrics (**Figure 3, Supplementary Table S2**, and **Supplementary Figures S1** and **S2**) and achieved a robust performance across the two naming schemes. For example, via the ‘protein complex’-leave-one-out cross-validation strategy, the RF classifier achieved AUC values of 0.885 and 0.895 for biological function, 0.925 and 0.927 for molecular function, and 0.951 and 0.957 for cellular component category for protein complexes with conventional and gene combinational names, respectively. In addition, we also observed that the resulting GO term lists predicted by the RF classifiers were able to significantly reduce the number of nearest neighbors needed to recover the majority of the ground-truth GO term annotations for a protein complex when compared to the nearest-neighbor results from querying the *k*-d tree (**Figure 3D**). For example, for protein complexes with subunit names, the RF classifier predicted terms were able to recover 80.5%, 83.6%, and 89.2% of CORUM’s ground-truth in 102, 49, and 11 positively predicted terms for biological process, molecular function, and cellular component, respectively.

**Figure 3.**
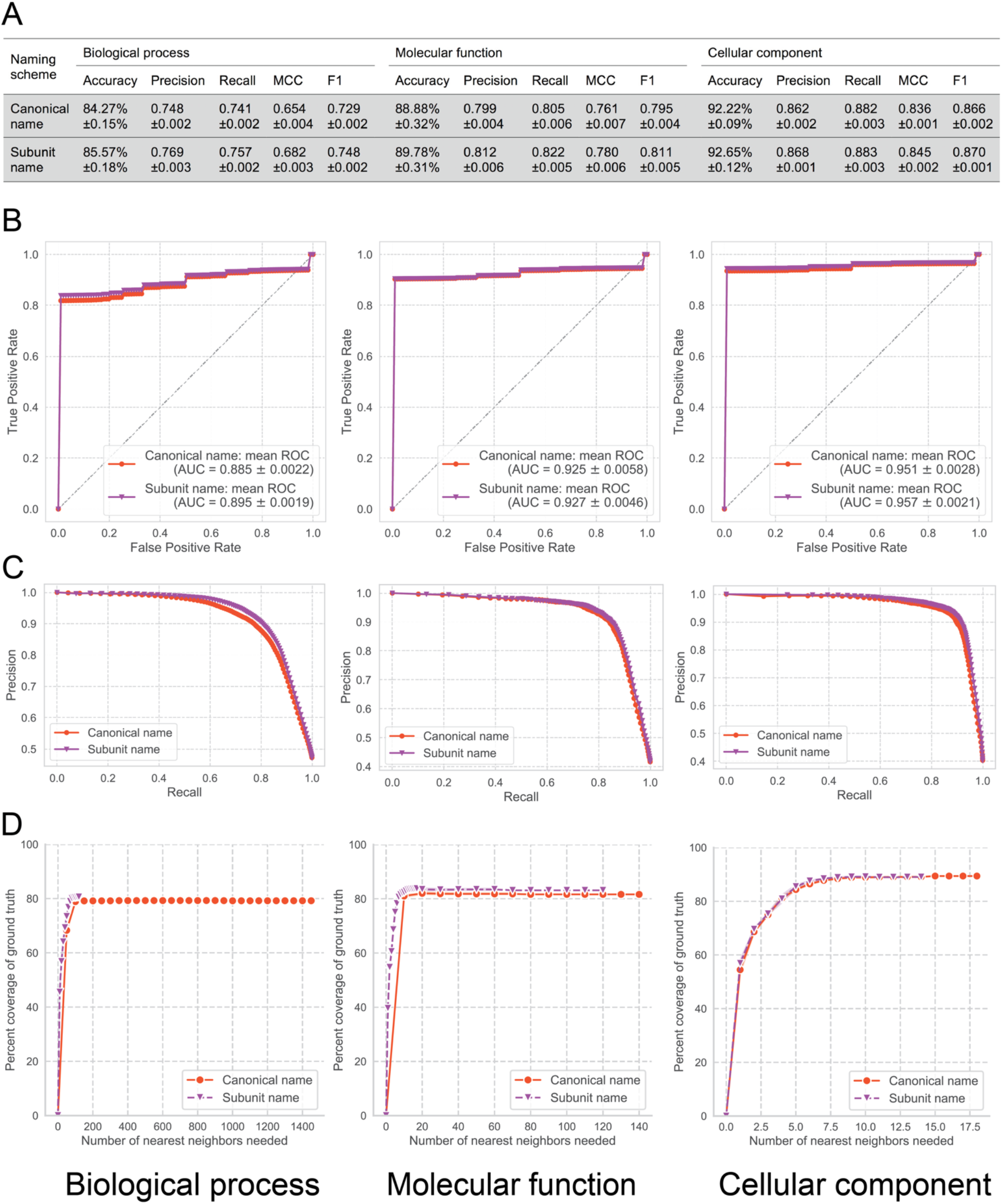
Prediction performance of RF model trained on the ground-truth protein complex – GO term annotations in the CORUM database for biological process, molecular function, and cellular component using canonical and subunit naming schemes for protein complexes, respectively, including (A) performance measures of RF models; (B) ROC curves and mean AUC values; (C) the precision-recall curves of the RF models via the adapted protein complex leave-one-out cross-validation, and (D) the numbers of predicted GO terms by the RF models to recover the CORUM database annotations.

While the RF classifiers performed well to recover the ground-truth as documented in the CORUM database, performance of the supervised approach may belie the inherent bias to the database that it was trained upon. Although protein complexes within CORUM have been extensively studied and the GO term annotations have been manually curated, there is an extant right skew in the frequency of GO terms with low to middle depth, based on the GO DAG structure. As shown in **Supplementary Figure S3**, we observed that the logged frequency of a particular GO term (i.e. the number of times a GO term has been annotated in CORUM) versus each GO term’s depth in the GO DAG structure reveals a biased annotation distribution for GO terms in CORUM. For example, the biological process term ‘Regulation of transcription DNA templated’, molecular function term ‘DNA binding’ and cellular component term ‘Nucleus’ were annotated in 233, 278, and 702 protein complexes respectively out of 3511 total protein complexes in the CORUM database. Such over-annotated GO terms could bias machine-learning algorithms in favor of selecting these highly abundant annotations. Therefore, to address the biases of the dataset that the RF classifier was trained upon, we supplemented the predicted terms from the RF classifier with the predicted nearest neighbors from the *k*-d tree. A graphical illustration of the combination of the RF and *k*-d tree prediction lists together as well as an example is shown in **Figure 4** (STAR Methods).

**Figure 4.**
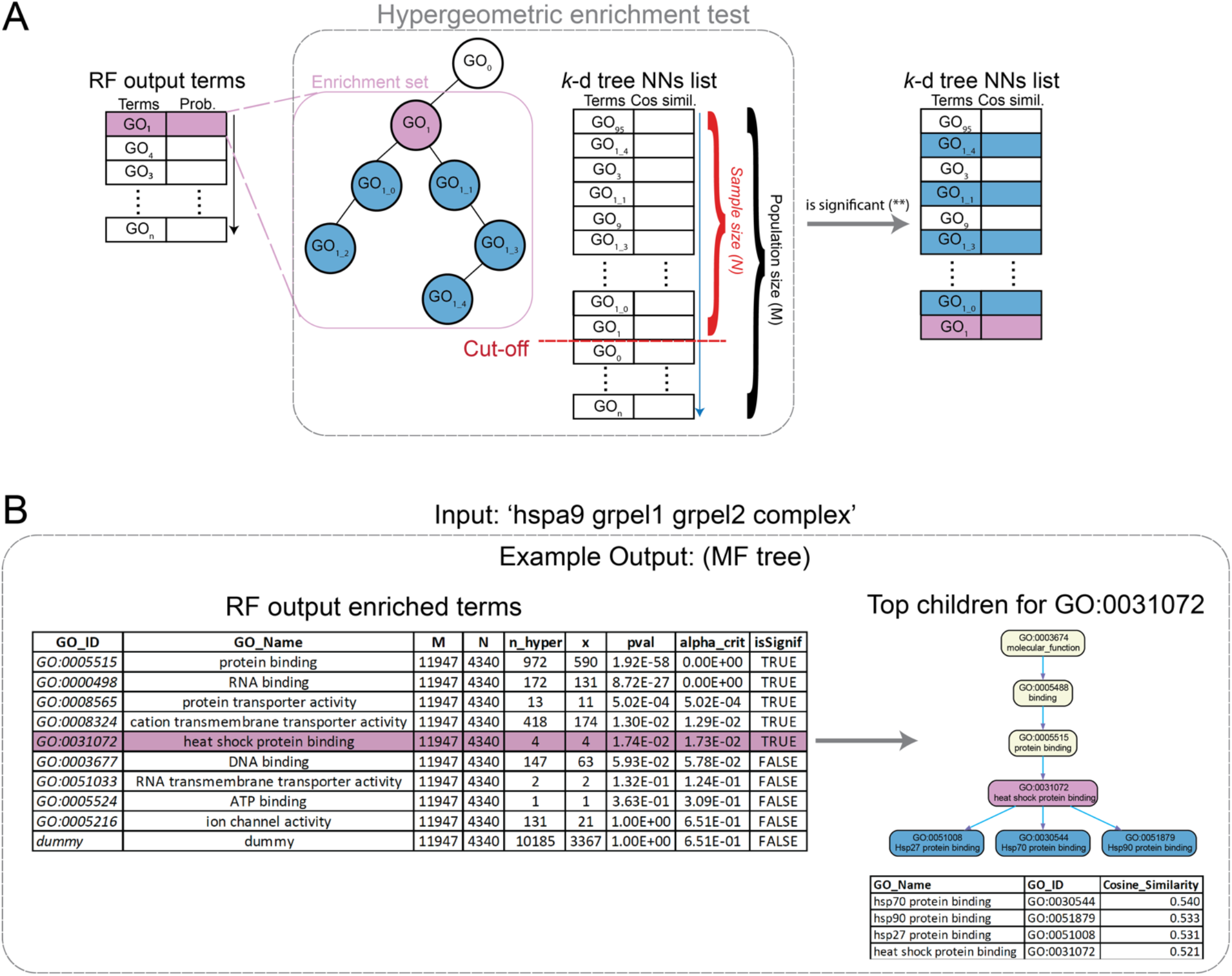
Utilizing functional enrichment analysis to combine the prediction lists from the *k*-d tree and the supervised RF model. (A) Hypergeometric enrichment test based on the RF prediction list. For each term in the list, all the child nodes of the term were collected and used for the statistical significance test with the terms from the *k*-d tree. If significant, the top ten terms from the *k*-d tree that are the child nodes of the RF term were selected and visualized in the GO DAG structure. (B) An example of prediction results for the protein complex ‘hspa9 grpel1 grpel2 complex’ illustrating the enrichment analysis procedure.

#### Independent test demonstrates divergent GO term predictions by PCfun compared to Complex Portal

We accessed the prediction performance of PCfun (trained on CORUM) with the testing data from a different database, Complex Portal. We first identified that only 110 annotated human protein complexes (i.e. complexes with identical subunits) were shared by CORUM and Complex Portal. We subsequently interrogated the semantic similarity of the biological process, molecular function and cellular component terms for these complexes between the two databases (STAR Methods). The heatmaps of the pairwise similarity scores using the method reported by Wang *et al* (Wang et al., 2007) are shown in the left panel of **Figure 5**. The average semantic similarity scores of biological process, molecular function and cellular component categories were 0.40, 0.34 and 0.54, respectively, suggesting that even for complexes with the same subunit composition the annotations of CORUM and Complex Portal are dissimilar. More stringently, we examined the numbers of identical GO terms used for each overlapping protein complex. As a result, less than one GO term across all categories (0.22 biological process term, 0.05 molecular function term and 0.13 cellular component term, respectively) was shared on average per protein complex between CORUM and Complex Portal. This means that the annotations for protein complexes are extremely divergent across the two databases, making it challenging for PCfun (built on CORUM) to accurately cover the GO annotations in Complex Portal. We then sought to gauge the approximate similarity of GO terms predicted by PCfun with Complex Portal annotations by assessing the pairwise semantic similarity (STAR Methods). The right panels of **Figure 5** show the comparison between predicted biological process, molecular function and cellular component terms by PCfun and the Complex Portal annotations for the non-overlapping protein complexes (i.e. with <50% of overlapping subunits). For biological process as shown in **Figure 5A**, 15 (approximately 44.1%) non-overlapping complexes covered 90-100% of similar terms (semantic similarity ≥ 0.5) compared to Complex Portal biological process annotations, while 14 complexes (41.2%) had divergent predictions (i.e. coverage between 0-10%) compared to the annotations in the Complex Portal. Similarly, for cellular component (**Figure 5C**), approximately 69.7% (23) of the complexes covered 90-100% of similar terms compared to Complex Portal cellular component annotations 30.3% (10). In contrast, PCfun demonstrated highly divergent predictions for the molecular function category with only 1 (3.2%) complex covering 90-100% of similar terms and 27 (87.1%) complexes demonstrating 0-10% coverage of similar terms when compared to Complex Portal’s annotations. The low coverage of PCfun prediction for the molecular function category might be related to the limited overlap between the molecular function annotations from CORUM and Complex Portal (the left panel of **Figure 5B**). It is noteworthy that different studies and databases may have divergent annotations for a protein or protein complex. Despite the low semantic similarities of all the biological process, molecular function and cellular component terms between the two databases, PCfun still demonstrates its ability to accurately recover and reliably predict functions accurately in the context of the database it was trained.

**Figure 5.**
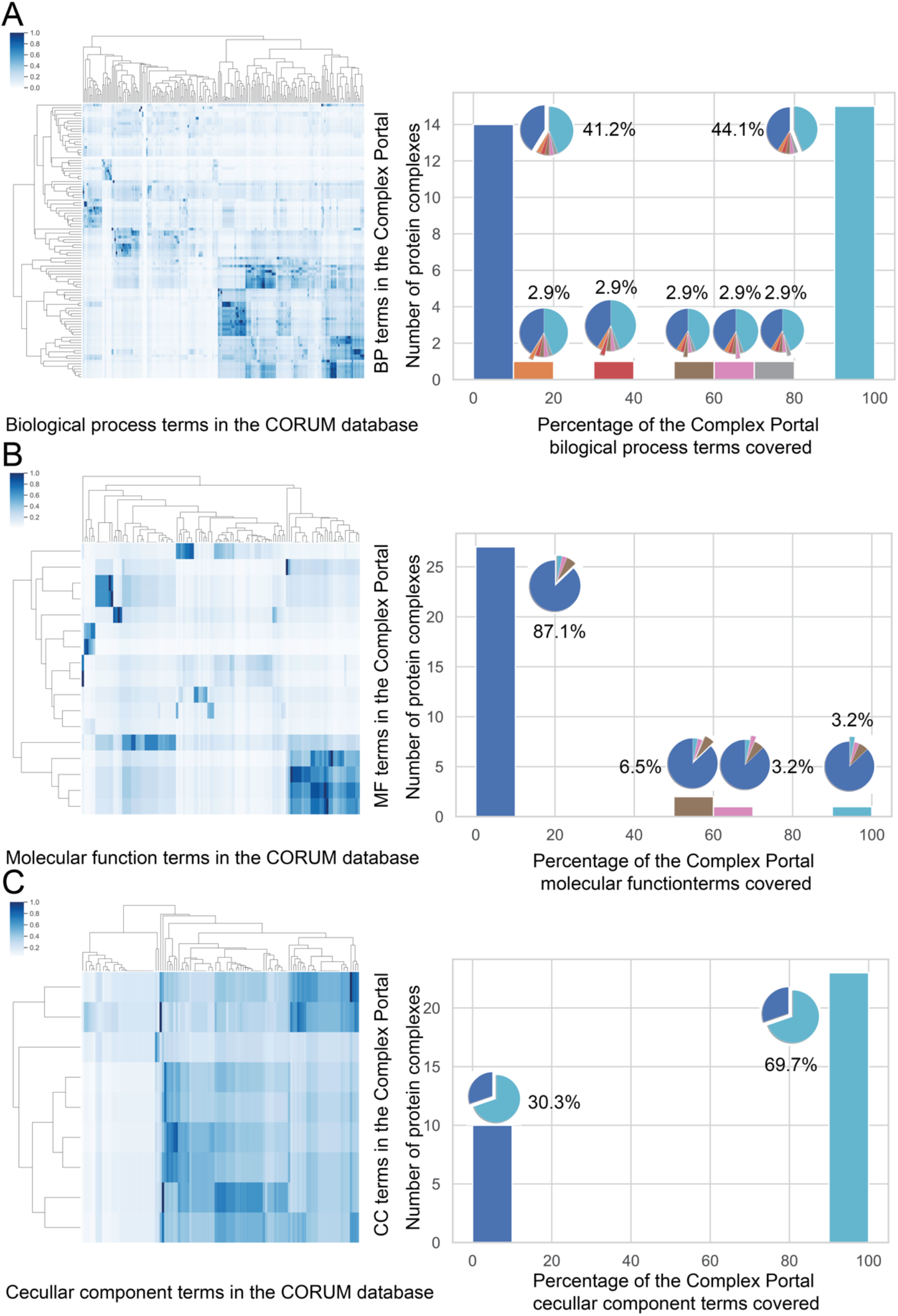
Semantic similarity comparison of (A) biological process, (B) molecular function, and (C) cellular component terms for the Complex Portal annotations with the CORUM annotations on identical protein complexes (left panel) and PCfun predictions on non-overlapping protein complexes (right panel). The right panel illustrates the percentage breakdowns of the predicted similar biological process and molecular function terms by PCfun to the Complex Portal annotations, respectively. Any GO term pairs between PCfun predictions and Complex Portal annotations with the semantic similarity ≥ 0.5 were considered similar.

## DISCUSSION

Advances in high throughput omics techniques have allowed for fast and deep identification and quantification of functional macro-molecules in the cell. Understanding the functions of these molecules, particularly for protein complexes, is a crucial step to understand and model biological activities in the cell. However, annotating the function of a protein complex is a great challenge, given the huge and occasionally contradictory volume of literature on each of its subunits. Currently, manual literature review is still the most common way to annotate the functions of protein complexes in major databases, such as CORUM and Complex Portal. Computational methods for gene function prediction (Barutcuoglu et al., 2006; Guan et al., 2008; Stojanova et al., 2013) have greatly broadened our understanding of single gene functions. However, combining the functional annotations of genes to infer the functions of the complex remains challenging and there is no such work published, to the best of our knowledge. In this study, we describe PCfun, the first hybrid computational framework for function prediction of protein complex, powered by the integration of machine-learning and text-mining techniques. PCfun was built based on large-scale publications obtained from PubMed Central (approximately one million full-text articles and their abstracts). The resulting word-embedding matrix was then used to build two complementary computational models, a supervised RF model for the prediction of function based on the annotations in the CORUM database, and an unsupervised *k*-d tree model for nearest-neighbor queries. The adapted protein complex leave-one-out cross-validation demonstrated the accurate prediction performance of the supervised RF model for predicting the PC-GO associations with respect to the ground-truth annotations provided in CORUM. On the other hand, we also constructed the *k*-d tree model, which is able to shortlist top-ranked nearest-neighbors, including terms not annotated in the CORUM database. Through enrichment analysis of the functional terms predicted by both models, PCfun is able to provide the final predicted GO terms associated with the input protein complex. Its representation of a contextual GO dendrogram structure along with the embedded predicted GO terms illustrates the hierarchal relationships of all predicted GO terms for the given protein complex.

As discussed in the ‘Results’ section, one issue during the construction of the machine-learning models is the biased functional annotations in the CORUM database, as shown in **Figure S3**. Therefore, the predictive power of the RF model in PCfun is limited to the CORUM annotations, demonstrating that it is crucial to combine the ‘non-biased’ prediction results of unsupervised *k*-d tree method with RF predictions via enrichment analysis, in order for PCfun to deliver non-biased predicted functions for a given protein complex. Another noteworthy issue is the negative data for training the supervised RF model of PCfun. Traditional supervised-learning models for binary classification require accurate labeling of classes (e.g. positive vs. negative) for each sample in the training dataset. However, labelling negative training samples is practically challenging in biological context due to the lack of experimental data. Oftentimes a previously labelled negative sample can be relabeled as positive with the acquisition of new data from novel biological techniques or identification methods. We will re-train the PCfun model once the annotations in the referred databases are updated, in order to continually improve the prediction performance.

PCfun can be applied to broad biological and personalized medicine applications. As demonstrated in this work, PCfun is easily accessible and offers utility to a variety of proteomic technologies. Prior to the prediction, the potential protein complexes with clearly defined subunits (either UniProt accessions or gene names) can be provided using the methodology described in the study of McBride *et al* (McBride et al., 2019). It is also possible to compare the functional differences by examining the prediction outputs of PCfun for a protein complex across different biological/medical conditions. In the future, we also plan to incorporate the differential analysis, similar to gene set enrichment analysis, for differentially regulated protein complexes that may have the ability to leverage both compositional (i.e. stoichiometric) and abundance differences. Additionally, prior to delivering the prediction results, PCfun first searches the given protein complex within the CORUM database for possible annotations. The documented annotations in CORUM and newly predicted GO terms are separated in the final outputs in an effort to facilitate the generation of novel hypotheses regarding the function of the protein complex. Taken together, we anticipate that PCfun can serve as an instrumental computational approach for the prediction of novel functions of protein complexes to provide reliable computational evidence for further experimental validation.

## ACKNOWLEDGEMENTS

This work was supported by in part by the Swiss National Science Foundation (grant No. 3100A0-688 107679 to R.A.) and the European Research Council (ERC-20140AdG 670821 to R.A.). C.L. is currently supported by a National Health and Medicine Research Council of Australia (NHMRC) CJ Martin Early Career Research Fellowship (1143366). M.B. was funded by an SNSF SystemsX.ch fellowship (TPdF 2013/135). A.W.P. is supported by an NHMRC Principal Research Fellowship (1137739). We thank Fabian Frommelt, Federico Uliana, Michiel Karrenbelt, Elias Pratschke and Shreyans Jain from ETH Zürich for their critical comments and insightful suggestions.

## AUTHOR CONTRIBUTIONS

R.A., C.L., M.M., V.S.S. and J.S. conceived and designed the project. V.S.S., C.L. and M.M. developed and implemented the PCfun package, and conducted data analysis, machine-learning prediction, and GO term enrichment analysis. M.M. assist with the generation of word-embedding and *k*-d tree. A.F. and M.M. assisted with package implementation and data analysis. R.C., E.G.W., M.B., W.S., P.G.A.P, M.R.M., Z.C. and A.W.P. provided critical and insightful comments during the development of PCfun. V.S.S., C.L., J.S. and R.A. drafted the manuscript, which has been revised and approved by all the other authors.

## STAR METHODS

### KEY RESOURCES TABLE

| REAGENT OR RESOURCE | SOURCE | IDENTIFIER |
| --- | --- | --- |
| Deposited Data |  |  |
| CORUM Database | (Giurgiu et al., 2019) | https://mips.helmholtz-muenchen.de/corum/ |
| Gene Ontology Resource | (Ashburner et al., 2000; The Gene Ontology, 2019) | http://geneontology.org/ |
| QuickGO database | (Binns et al., 2009) | https://www.ebi.ac.uk/QuickGO/ |
| Complex Portal | (Meldal et al., 2019) | https://www.ebi.ac.uk/complexportal/home |
| PubMed Central | - | https://www.ncbi.nlm.nih.gov/pmc/ |
| UniProt Database | (UniProt, 2019) | https://www.uniprot.org/ |
| Software and Algorithms |  |  |
| Scikit-learn | (Pedregosa et al., 2011) | https://scikit-learn.org/stable/ |
| fastText | (Joulin et al., 2017) | https://fasttext.cc/ |
| PCfun | This work | https://github.com/sharmavaruns/PCfun |

### LEAD CONTACT AND MATERIALS AVAILABILITY

Further information and requests for resources should be directed to and will be fulfilled by the Lead Contact, Chen Li.

## METHOD DETAILS

### Text corpus generation and data processing

Approximately 1 million articles (including open-access full-text articles and their abstracts) were downloaded from PubMed Central in February 2018. Note that these publications are not species/organism specific, which means that the developed PCfun, built on the corpus, is a generic tool for protein complex function prediction. For processing the articles into a text corpus, we followed the text processing pipeline described in the study of Manica *et al* (Manica et al., 2019). All of the natural language queries were pre-processed by removing all punctuation characters, by fixing Unicode mojibake and garbled HTML entities, and by converting all uppercase characters into lowercase. For the extraction of a single word vector from a natural language query (e.g. a protein complex or GO term name), L2 normalized vectors of the query individual words were extracted from the embedding and averaged. The final averaged vector of the component vectors of the name were L2 normalized again and subsequently used as the final word vector for the natural language query.

### Word-embedding and similarity calculations

Word-embedding refers to a class of approaches developed in the fields of text mining and natural language processing that embed natural language texts into high-dimensional, continuous real-valued vector representations (Manica et al., 2019). The word-embedding in this study was achieved by training the unsupervised version of ‘fastText’ (Joulin et al., 2017) with a skip-gram model on the publications corpus. Briefly, the skip-gram model attempts to minimize the negative sum of the log probability that a word exists within the context of a target word. The fastText unsupervised training parameters used were the default values as declared in the fastText package except for 500-dimensions for the embedding layer, a context window of size 9, and the usage of bi-grams as chosen from the study of Manica *et al* (Manica et al., 2019). After training, the word vectors were normalized to a unitary norm. We utilized the cosine similarity between two vectors, which measures the cosine of the angle between the two vectors as the normalized dot product of the two vectors.

### Calculating sub-embeddings of topics and nearest neighbors

We extracted sub-embeddings (a data frame consisting of the word vectors for a particular class of terms, such as all biological process GO terms) for protein complex names (extracted from CORUM- one sub-embedding for each naming scheme: *canonical* or *subunit* names) and GO terms (extracted from GO resource split into biological process, molecular function and cellular component classes). For example, to obtain the GO terms most similar to a protein complex query in the embedding space, we calculated of the nearest neighbors to the protein complex query vector within the extracted GO term sub-embedding space. A nearest-neighbor calculation involves calculating distances between vectors within the sub-embedding of interest and the query vector, and then sorting the neighbors by their similarity by descending order. The calculation for nearest neighbors can be quite time intensive depending on the size of the embedding due to the time requirements for pairwise calculation of the distances between each vector in the sub-embedding and the query vector. Therefore, we stored the sub-embedding vectors into a pre-computed *k*-d tree (*k*-dimensional tree), which is a space-partitioning data structure that stores the vectors into buckets determined by hyperplane splits over each dimension of the vector (Freidman et al., 1977). Therefore, calculation of *n*-nearest neighbors to a query vector requires only placement of the query vector into its corresponding location within the pre-calculated *k*-d tree and then subsequent querying of its ancestors in the tree until the closest *n*-nearest neighbors have been calculated.

### Databases for protein complex annotations

For this work we employed the CORUM database (Giurgiu et al., 2019) as the main resource for the ground-truth annotation of protein complexes with GO terms, as CORUM is a compendium of manually curated and experimentally validated protein complexes for various organisms (Giurgiu et al., 2019). Annotations in CORUM for the function of protein complexes have been collected from various types of evidence, including experimental evidence (‘exp’), evidence from literature (‘lit’), known mammalian homologs (‘kmh’), high-throughput experiments (‘htp’), and predicted function (‘pred’). Here the ‘predicted function’ refers to the potential function suggested by the experimental results. In this work, we utilized annotations from all species in the CORUM database to keep as much information as possible for constructing an accurate supervised machine-learning model, given that the corpus we obtained from PubMed are not species/organism specific. In our study, non-redundant 3414 core protein complexes (downloaded in March 2019) from the CORUM database were used.

In addition to CORUM, we utilized the annotated *Homo sapiens* protein complexes (downloaded in November 2019) from the Complex Portal database (Meldal et al., 2019) to independently assess the prediction performance of PCfun. Similar to CORUM, the Complex Portal contains the protein complexes and their annotations of GO terms. As the Complex Portal has fewer number of protein complexes documented, we did not use it for training the model. For the independent test, only the protein complexes annotated as ‘physical interaction evidence used in manual assertion’ with the evidence code ‘ECO:0000353’ coupled with experimental evidence from the IntAct database (Orchard et al., 2014) were retained. To objectively benchmark the performance of PCfun on the Complex Portal, we further removed those protein complexes from the Complex Portal that had a subunit overlap of larger than 50% compared to the complexes in the CORUM database. As a result, the numbers of protein complexes from the Complex Portal for the independent test were in total 34, of which 34, 31, and 33 protein complexes have biological process, molecular function and cellular component annotations, respectively.

### Gene name extraction for protein complex subunits

As each unique natural language query has its own unique word vector, it is important to standardize the natural language name used for each protein complex when extracting its word vector. A protein complex can be either represented as its documented name, as written into the protein complex database, or as its subunits’ gene names strung together. In this study, we tested the performance of the algorithms using two different naming schemes: (1) *canonical name* (as documented in CORUM), and (2) *subunit name* (composed by UniProt gene names of each subunit). To obtain the gene names of the subunits, we extracted the subunits of the protein complexes from CORUM and queried the UniProt database (downloaded in May 2019) (UniProt, 2019) for their corresponding gene names using the organism identifier from the CORUM database. We then represented the protein complex name with its text pre-processed subunits’ gene names, strung together with spaces demarcating each individual gene name. Due to the fact that there might exist multiple names for a single gene, only its canonical name was extracted and used in our model. Gene names were extracted by downloading the UniProt FASTA-formatted sequence file, with respect to the appropriate species, which was then subsequently parsed for each relevant UniProt ID - gene name pair. The resulting work has tested the performance of the algorithm when using each naming scheme independently from each other. Importantly, use of the subunit UniProt gene name scheme also allows for greater flexibility as one can still gain functional insight into a newly detected protein complex even if the complex has not been officially named yet.

### Preparation of protein complex - GO (PC-GO) pair datasets for supervised learning

To enable the accurate prediction of protein complex function annotations, we have formulated our task as a supervised binary classification problem, for which we created a labelled dataset based on the CORUM annotations. To create the labelled dataset, we first extracted PC-GO term pairs and then labelled each pair as positive if its annotation was observed in the CORUM database. The GO terms and their DAG structures were collected from the Gene Ontology Resources platform (Ashburner et al., 2000; The Gene Ontology, 2019). To label the negatives, we first generated a pool of all possible negative PC-GO pairs to sample from by taking the GO terms (split by biological process, molecular function and cellular component categories) that were used in CORUM and not annotated for a particular protein complex. Since only a few GO terms were annotated for each protein complex, this negative sample pool significantly outnumbered the positive sample pool. This issue of having a huge number of negatives compared to positive labels is a common problem shared by a variety of bioinformatics studies and have been discussed in the recent studies of Li *et al* (Li et al., 2018b; Li et al., 2019a). To build an unbiased supervised classifier, it is common practice to train on an approximately equal distribution of positive and negative labels. To ensure this, we randomly selected an approximately equal number of negative PC-GO pairs from the pool of negatives as there were positives for each protein complex. This random selection was repeated five times, resulting in five training datasets, where only negative samples were different.

### Supervised machine-learning classifiers

Supervised binary classification was performed with the RF (Breiman, 2001), LR (Lecessie and Vanhouwelingen, 1992; Yu et al., 2011), and NB (with Gaussian and Bernoulli distributions, respectively) (Zhang, 2004) classifiers with the default parameters. The feature space consisted of 1,000-dimensional vectors (500-dimensional protein complex vector prepended to a 500-dimensional GO term vector) with each vector corresponding to a PC-GO pair labelled with either positive or negative. *RF* classification uses the ensemble of decision trees that randomly bootstraps over the training data and features for each decision tree in order to classify the input vector space. After construction of the random decision trees, the classifier outputs a class membership (positive or negative for binary classification) prediction dependent on the community-wide vote from the random trees constructed. *LR* utilizes the logistic function for the binary classification of a dependent variable. This method outputs a probability score ranging from 0 to 1 where values of >0.5 are considered to be of the positive class membership. *NB classifiers* are probabilistic classifiers built upon Bayesian statistics. These classifiers attempt to learn the distributional fit for each labelled class and accordingly assign probability values for each class when given a new input vector. The priors we tested were the Gaussian and Bernoulli distributions. Briefly, the Gaussian distribution assumes that the continuous data values are distributed according to a normal distribution, whereas the Bernoulli distribution assumes that the features are independent binary variables and can be considered as a special case of the binomial distribution. These two classifiers are termed NB_Gauss and NB_Bernoulli in the following sections, respectively. For PCfun’s machine learning classifier predicted terms we used a majority voting scheme over the five datasets to provide an averaged predicted probability for each GO term. For this majority vote, we equally weighted the contribution of each dataset’s model to achieve a final combined probability score for particular GO terms. If the combined probability score was >0.5, the GO term was classified as a positive term, otherwise classified as a negative.

### Performance assessment of supervised binary classification

To assess the predictive performance of the supervised binary classifiers, we introduced an adaptation on traditional leave-one-out cross-validation, which we termed ‘protein complex’-leave-one-out cross validation. The ‘protein complex’-leave-one-out cross validation first pre-removed every row that had the particular protein complex being tested (including both positive and negatively labelled rows). Afterward, the model was then trained on the remaining dataset and tested on the pre-removed protein complex of interest’s rows. This complex-wise evaluation strategy was applied to each protein complex in the dataset. To measure the performance, we used widely established performance metrics used in a variety of bioinformatics and computational biology studies (Li et al., 2018a; Manavalan et al., 2019; Song et al., 2018), including accuracy, AUC (Area Under the Curve), precision, recall, MCC (Matthews Correlation Coefficient) (Matthews, 1975), and F1 score. All metrics reported in this study are the average of five datasets of ‘protein complex’-leave-one-out cross-validation. For plotting the Receiver Operating Characteristic (ROC) curves, we followed the recommendations of the interpolation scikit-learn package (please refer to the scikit-learn tutorial). These scores are calculated from the elementary scores of true positives (*TP*), true negatives (*TN*), false positives (*FP*), and false negatives (*FN*). The formulas for each performance metric are shown as follows:

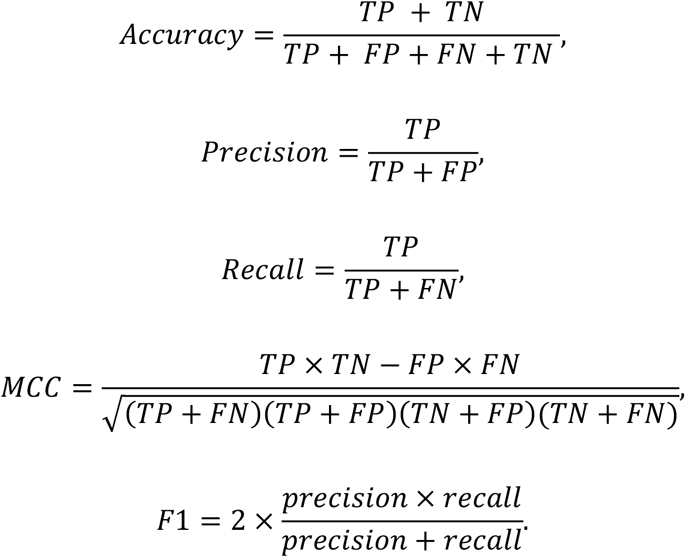

### Semantic similarity calculation between GO terms

To compare the GO terms predicted by PCfun and the annotations from Complex Portal, we did not perform the direct comparison/matching of the GO terms, due to the low number of intersecting GO terms between CORUM and Complex Portal. Instead, we exploited the semantic similarity introduced by Wang *et al* (Wang et al., 2007). This method considers both biological meanings and hierarchical relationships in the GO DAG structure of the given GO term pair. Any GO term pairs with a semantic similarity ≥0.5 were considered similar. When comparing the predicted GO terms by PCfun and the annotations from Complex Portal, for each protein complex, we first calculated the similarities for all possible GO term pairs between the PCfun outputs and Complex Portal annotations. The pairs with similarity score ≥ 0.5 were selected and the unique Complex Portal GO terms (*N*) in the pairs counted. Then the percent coverage of the predicted GO terms to the Complex Portal annotation for the particular protein complex was calculated using *Coverage* = (*N*/*M*) × 100, where *M* denotes the number of GO terms annotated in Complex Portal for the particular protein complex.

### GO term enrichment analysis of predicted functional terms

This step aims to systematically and comprehensively combine the two predicted GO term lists in each category (i.e. biological process, molecular function, and cellular component) by RF and the *k*-d tree for a given protein complex, respectively, due to the fact that this shortlist of terms predicted by RF that recovers CORUM database well but tends to predict broader GO terms for a protein complex. Given the predicted GO term list by RF with the size *N* (*N*≤10), we supplemented the information from this list with the list of GO terms (i.e. the nearest neighbors) obtained directly by querying the GO term sub-embedding. To accomplish this, we developed a functional enrichment analysis pipeline based on the hypergeometric test to assess if all the child nodes of the GO term by RF are significantly enriched in the predicted GO terms by the *k*-d tree, using the following formula:

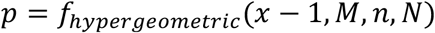

where *M* denotes the number of total GO terms for a particular GO term class, *n* denotes the number of child terms of a parent GO term plus the parent term predicted by the supervised classifier that exists within the specific GO term class, where *N* denotes the sample size which is the mean number of GO terms required for the nearest neighbor list to recover all GO terms annotated for a protein complex (biological process = 11044, molecular function = 5213, and cellular component = 1896, respectively), and *x* denotes the number of child terms for a particular supervised term that exist in the set of the sample size list. The function *f*_*hypergeometric*_ was the survival function for the hypergeometric distribution as implemented in the SciPy package. If the child nodes/terms are significantly enriched in the predicted list by the *k*-d tree, 10 top ranked terms based on cosine similarity from the *k*-d tree list are selected. We could therefore obtain a ‘combined’ predicted list that not only accurately recovers CORUM database but also supplements the list by RF using the detailed GO terms predicted by the *k*-d tree. To visualize the results, PCfun plots a GO tree structure of predicted GO term by RF (in green) and the 10 top ranked GO terms by the *k*-d tree (in purple) to demonstrate the hierarchical relationships of these terms. For the cellular component category, in addition to the combination of the two lists from the k-d tree and the RF model by functional enrichment analysis, we also considered adding the overlap of cellular component annotations of all the subunits to the final outputs. The cellular component annotations for each subunit was downloaded from the QuickGO database (Binns et al., 2009).

### PCfun prediction output organization

In total, there are six output lists (two for each GO category) for a given protein complex generated by PCfun. For each GO category (i.e. biological process, molecular function, and cellular component), one list contained the RF predictions and the top 10 significantly enriched GO terms by the *k*-d tree, while the other lists provided the top 20 GO terms by the *k*-d tree only. In addition, for each RF predicted term, a GO DAG structure is plotted to illustrate the hierarchical relationships between the RF prediction and the top 10 significantly enriched terms from the *k*-d tree.

## DATA AND CODE AVAILABILITY

### Data and code availability statement

#### Data availability

The full-text articles and their abstracts (in the non-commercial use collection) were extracted from the PubMed Central under a Creative Commons or similar license. The training dataset and the validation test (i.e. the PC-GO association) were obtained from the CORUM database (Giurgiu et al., 2019) and the Complex Portal database (Meldal et al., 2019), respectively. The full GO lists were downloaded from the Gene Ontology Resource platform (Ashburner et al., 2000; The Gene Ontology, 2019). For evaluation purposes, we downloaded the GO annotations for individual proteins from the QuickGO database (Binns et al., 2009).

#### Code availability

PCfun is an open-access software and is freely available for academic use under the ‘Academic Free License v3.0’. The source code, user instruction, and example inputs can be downloaded from https://github.com/sharmavaruns/PCfun.

## Supplementary information

**Table S1.** Top 10 GO terms shortlisted by the *k*-d tree for the protein complex “SMAD2-SMAD4-FAST1-TGIF-HDAC1 complex, TGF(beta) induced”.

| Number | GO term | Cosine distance | Cosine similarity | GO ID |
| --- | --- | --- | --- | --- |
| Biological process |  |  |  |  |
| 1 | jun phosphorylation | 0.90142032 | 0.52592264 | GO:0007258 |
| 2 | common partner smad protein phosphorylation | 0.91515871 | 0.52214994 | GO:0007182 |
| 3 | smad protein signal transduction | 0.9206383 | 0.52066024 | GO:0060395 |
| 4 | pathway restricted smad protein phosphorylation | 0.92724758 | 0.51887469 | GO:0060389 |
| 5 | regulation of histone h3 t3 phosphorylation | 0.93482421 | 0.51684282 | GO:2000281 |
| 6 | regulation of smad protein signal transduction | 0.93509793 | 0.51676971 | GO:0007184 |
| 7 | histone h3 t3 phosphorylation | 0.93731143 | 0.51617927 | GO:0072355 |
| 8 | negative regulation of smad protein signal transduction | 0.94274921 | 0.51473448 | GO:0060392 |
| 9 | regulation of pathway restricted smad protein phosphorylation | 0.9430544 | 0.51465363 | GO:0060393 |
| 10 | negative regulation of histone h3 k27 trimethylation | 0.94409184 | 0.51437899 | GO:1902465 |
| Molecular function |  |  |  |  |
| 1 | smad binding | 0.81312028 | 0.55153539 | GO:0046332 |
| 2 | i smad binding | 0.88015558 | 0.53187088 | GO:0070411 |
| 3 | r smad binding | 0.89042146 | 0.52898257 | GO:0070412 |
| 4 | co smad binding | 0.89897215 | 0.52660067 | GO:0070410 |
| 5 | bmp (bone morphogenic protein) binding | 0.93230116 | 0.51751767 | GO:0036122 |
| 6 | activin binding | 0.95586228 | 0.51128344 | GO:0048185 |
| 7 | histone deacetylase activity h3 k14 specific | 0.96414003 | 0.50912867 | GO:0031078 |
| 8 | bmp receptor binding | 0.96599249 | 0.50864894 | GO:0070700 |
| 9 | transcription corepressor binding | 0.97184839 | 0.50713838 | GO:0001222 |
| 10 | bmp receptor activity | 0.97334295 | 0.50675429 | GO:0098821 |
| Cellular component |  |  |  |  |
| 1 | pdx1 pbx1b mrg1 complex | 0.84789228 | 0.54115709 | GO:0034978 |
| 2 | gata2 tal1 tcf3 lmo2 complex | 0.86410241 | 0.53645121 | GO:0070354 |
| 3 | rgs6 dnmt1 dmap1 complex | 0.86604966 | 0.53589142 | GO:0070313 |
| 4 | gata1 tal1 tcf3 lmo2 complex | 0.87113766 | 0.53443422 | GO:0070353 |
| 5 | heteromeric smad protein complex | 0.8795976 | 0.53202877 | GO:0071145 |
| 6 | smad protein complex | 0.88390432 | 0.53081252 | GO:0071141 |
| 7 | maml3 rbp jkappa icn1 complex | 0.89328291 | 0.52818308 | GO:0071179 |
| 8 | maml1 rbp jkappa icn1 complex | 0.89389121 | 0.52801343 | GO:0002193 |
| 9 | homomeric smad protein complex | 0.90011405 | 0.5262842 | GO:0071143 |
| 10 | fhl2 creb complex | 0.90109935 | 0.52601144 | GO:0034980 |

**Table S2.**
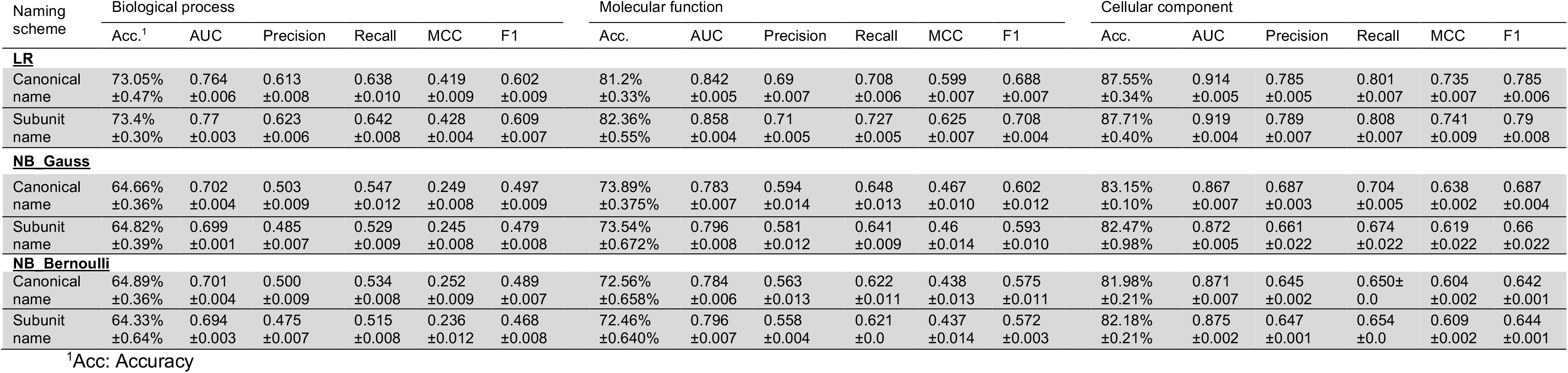
Performance comparison of the Logistic Regression (LR) and Naïve Bayes (NB_Gauss and NB_Bernoulli) classifiers trained on the CORUM database via the adapted protein-complex leave-one-out cross-validation, for biological process, molecular function and cellular component categories, respectively.

**Figure S1.**
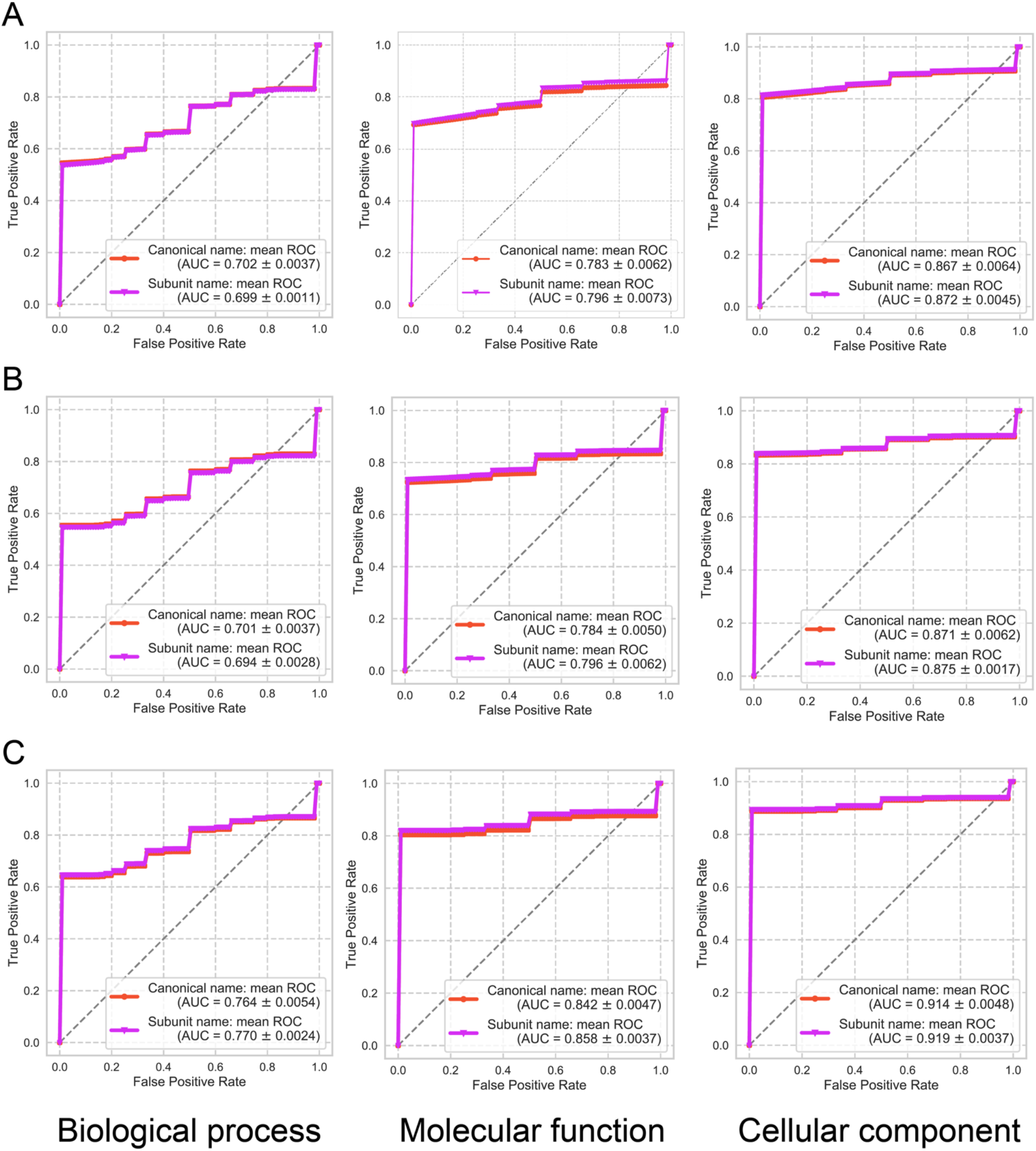
ROC curves and the average AUC values of (A) NB_Gauss, (B) NB_Bernoulli and (C) LR models on biological process, molecular function and cellular component categories via the adapted protein complex leave-one-out cross-validation. These models were trained using the CORUM ground-truth PC-GO associations, where the protein complexes were represented using both canonical and subunit naming schemes.

**Figure S2.**
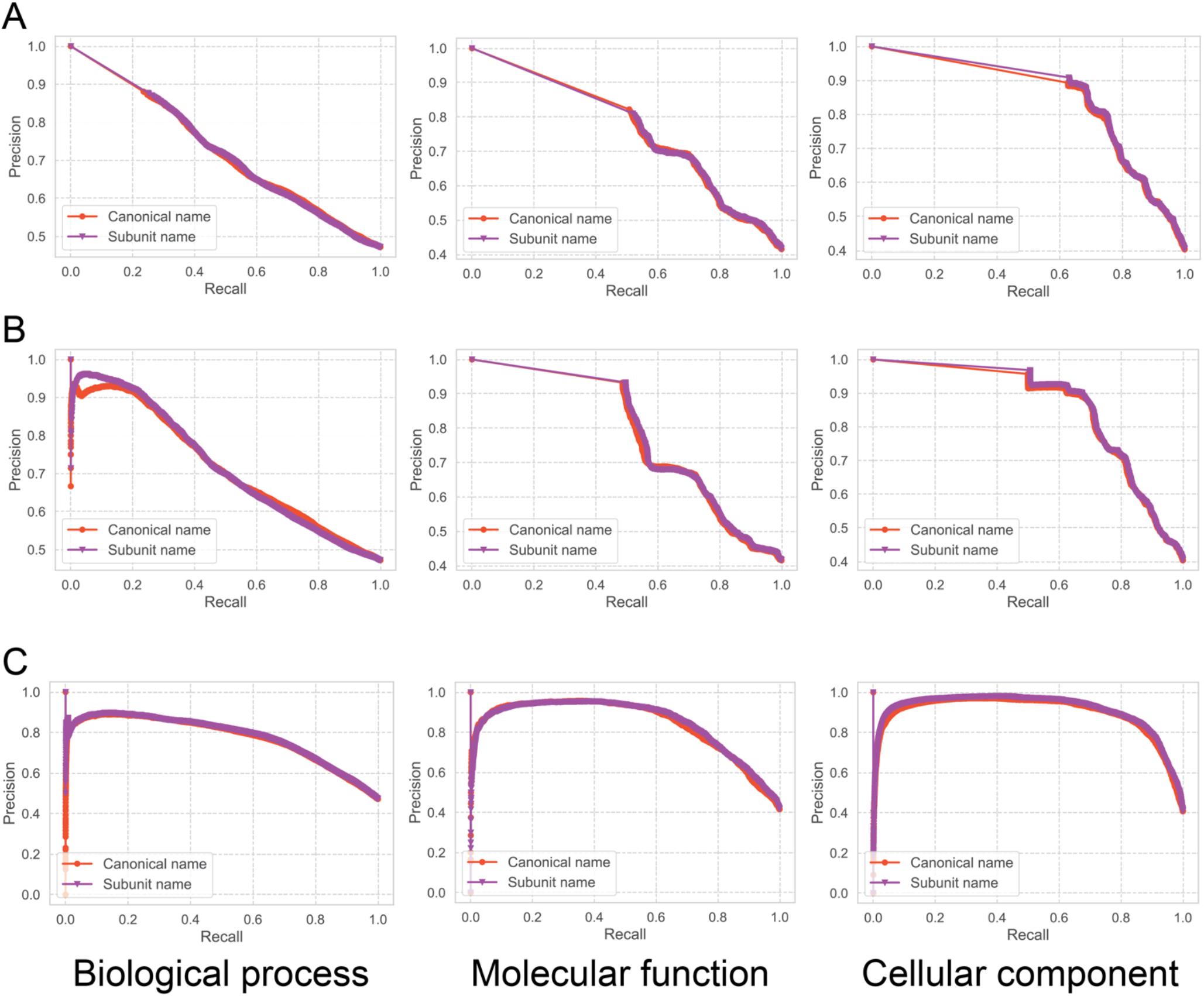
Precision-recall curves of (A) NB_Gauss, (B) NB_Bernoulli and (C) LR models on biological process, molecular function and cellular component categories via the adapted protein complex leave-one-out cross-validation, where the protein complexes were represented using both canonical and subunit naming schemes.

**Figure S3.**
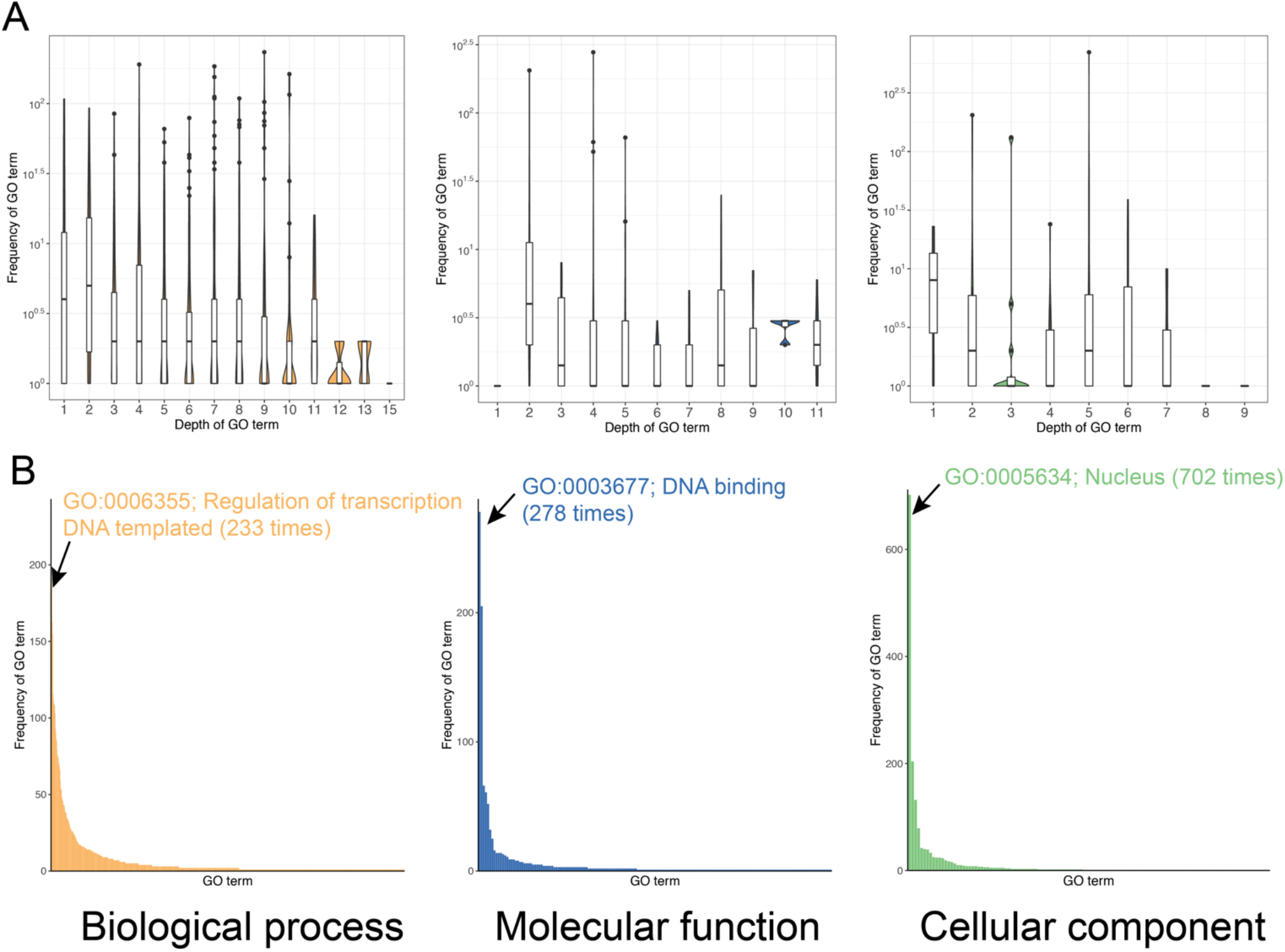
Statistical analyses of the annotated GO terms in the CORUM database, including (A) the distributions of depth of annotated biological process, molecular function and cellular component terms in the GO DAG structures and (B) the frequencies of biological process, molecular function and cellular component terms that were assigned to the CORUM protein complexes. The top over-annotated biological process, molecular function and cellular component terms are respectively indicated.

